# Bidirectional Crosstalk Between Bladder Cancer Cells and Normal Fibroblasts Drives Phenotypic Reprogramming and Mitomycin C Resistance

**DOI:** 10.1101/2025.09.09.675061

**Authors:** Jinhui Gao, Cindy Xinyu Ji, Ci Ren, John Hines, Eleanor Stride, Richard T. Bryan, Jennifer L. Rohn

**Affiliations:** Division of Medicine, University College London, London, UK; Cancer Research UK Manchester Institute, University of Manchester, Manchester, UK; Westmoreland Street University College London Hospital, London, UK; Institute of Biomedical Engineering, Department of Engineering Science, University of Oxford, Oxford, UK; Bladder Cancer Research Centre, Department of Cancer and Genomic Sciences, College of Medicine and Health, University of Birmingham, Birmingham, UK

**Keywords:** Bladder Cancer, Normal fibroblast, Cancer-associated fibroblast, Tumour microenvironment, Mitomycin C

## Abstract

During early bladder cancer progression, invading tumour cells first encounter normal fibroblasts (NFs) residing in the lamina propria. Whether and how this interaction shapes tumour behaviour and treatment response, however, has not been systematically examined. While cancer-associated fibroblasts (CAFs) are established drivers of tumour progression and therapy resistance, the role of their precursor NFs in shaping bladder cancer behaviour has been largely overlooked. Here we dissect bidirectional interactions between bladder cancer cells and NFs using indirect (conditioned media) and direct co-culture systems. Fibroblast-conditioned media (FCM) significantly reduced proliferation while accelerating migration in both RT112 and T24 bladder cancer cells, consistent with a pro-migratory “go-or-grow” phenotypic shift. Immunofluorescence revealed FCM-induced cadherin switching reduced E-cadherin and elevated N-cadherin, indicative of an epithelial-to-mesenchymal transition (EMT)-like state. Reciprocally, NFs acquired CAF-like features within 48 hours of tumour cell exposure, with significant upregulation of αSMA and FAP confirmed by both immunofluorescence and flow cytometry. Functionally, increasing fibroblast-to-cancer cell ratio progressively attenuated mitomycin C (MMC)-induced cytotoxicity, demonstrating a stromal-mediated, ratio-dependent protective effect against intravesical chemotherapy. Data from the Cancer Genome Atlas (TCGA) for Urothelial Bladder Carcinoma provided corroboratory evidence that fibroblast-enriched tumours exhibited a strong transcriptional correlation with EMT, elevated resistance-associated transcriptional signatures, and significantly poorer overall survival. Collectively, these findings establish tumour-NF crosstalk as a critical and bidirectional regulator of bladder cancer phenotypic plasticity and chemoresistance, and suggest that fibroblast-targeting strategies combined with intravesical chemotherapy may offer a rational approach to overcoming stromal-mediated treatment failure.

## Introduction

Bladder cancer is among the most common malignancies worldwide, with non-muscle-invasive disease (NMIBC) accounting for 75% of diagnoses [1]. Despite standard-of-care transurethral resection, BCG immunotherapy and intravesical mitomycin C (MMC), recurrence rates reach 50–80%, and 10–20% of cases progress to muscle-invasive disease [2,3]. The limited and variable efficacy of MMC points to microenvironmental factors as critical modulators of treatment response [4], yet the stromal determinants of resistance remain poorly defined.

The tumour microenvironment (TME), particularly its fibroblast compartment, is an established driver of cancer progression and therapy resistance [5,6]. Cancer-associated fibroblasts (CAFs) promote tumour invasion, ECM remodelling, and immune evasion through paracrine and contact-dependent mechanisms [7,8]. CAFs can arise from multiple precursors, most prominently NFs [9]. NFs are the predominant stromal cell type residing within the lamina propria, a layer of connective tissue composed of ECM, fibroblasts, endothelial cells, smooth muscle cells, and immune cells that lies immediately beneath the urothelium [10]. This anatomical position is particularly significant: as bladder cancer progresses from the urothelium (NMIBC) through the lamina propria toward the detrusor muscle (MIBC), invading tumour cells inevitably encounter NFs before any other stromal component, making NFs the first responders in the tumour microenvironment [10]. Yet despite their likely role as CAF precursors and their strategic position along the invasion axis, NFs remain comparatively under-characterised in bladder cancer.

Studies in lung, colorectal, and pancreatic cancer demonstrate that NFs actively remodel cancer phenotypes through paracrine signals and ECM interactions [10–12], but equivalent bidirectional interactions remain uncharacterised in bladder cancer. Two processes are particularly underexplored: EMT-like reprogramming of cancer cells toward migratory states, and reciprocal CAF-like activation of NFs, marked by α-Smooth Muscle Actin (αSMA) and Fibroblast Activation Protein (FAP) upregulation. To the best of our knowledge, these processes have not been examined together as components of a coordinated tumour-stromal dialogue in bladder cancer, nor has their combined consequence for intravesical chemotherapy response been addressed.

We hypothesised that NFs actively reprogramme bladder cancer cells toward a migratory, EMT-like phenotype, while tumour cells reciprocally activate NFs toward CAF-like states that diminish MMC chemosensitivity. To test this, we employed indirect and direct co-culture models, characterised bidirectional phenotypic changes, assessed functional MMC resistance, and compared the results against data from the Cancer Genome Atlas for Urothelial Bladder Carcinoma (TCGA-BLCA) to establish clinical relevance.

## Results

### Fibroblast- and tumour-derived signals reciprocally suppress cell proliferation

To explore the bidirectional effects between cancer cells and NFs, we first assessed how fibroblast conditioned media (FCM) influenced cancer cell proliferation, and subsequently assessed how cancer cell-conditioned media affected NF proliferation. For the former, MTS assays were conducted on RT112 and T24 bladder cancer cell lines. FCM treatment significantly reduced the proliferation of RT112 and T24 cells in 48 hours or 96 hours (Figure 1a, 1b). The reciprocal interaction was equally pronounced: when NFs were treated with RT112-conditioned media (RCM) or T24-conditioned media (TCM), proliferation was significantly suppressed at 48 and 96 hours. RCM reduced proliferation to 67.5% at 48 hours and 62.2% at 96 hours, while TCM led to reductions of 70.0% and 68.2%, respectively, relative to untreated controls (Figure 1c, 1d).

**Figure 1:**
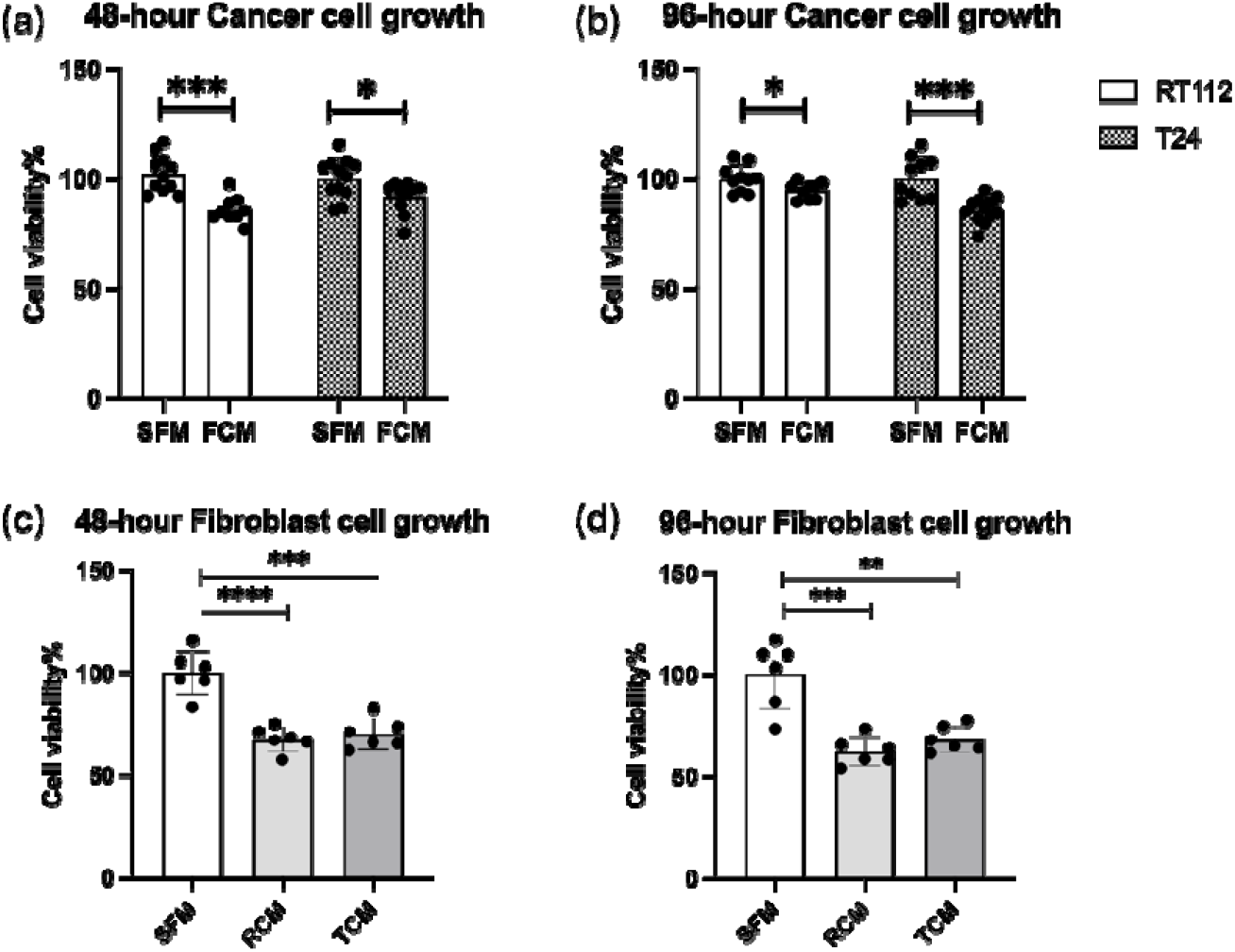
(a, b) Bar charts showing the proliferation of RT112 (low-grade) and T24 (high-grade) bladder cancer cell lines after 48 and 96 hours of treatment with FCM compared with serum-free DMEM (SFM). (c, d) Bar charts showing the proliferation of NFs at 48 and 96 hours following exposure to bladder cancer cell-conditioned media derived from RT112 (RCM) and T24 (TCM) compared with SFM.

### Fibroblast-derived signals promote EMT-like phenotypic plasticity and migration in bladder cancer cells

We investigated the impact of fibroblast-derived signals on bladder cancer cell behaviour using FCM. Exposure to NF-conditioned media significantly enhanced migration in both RT112 and T24 cells, as assessed by wound healing assays. In RT112 cells, FCM treatment consistently accelerated wound closure throughout the time course, producing a clear distinction between the two conditions. A similar effect was observed in T24 cells, where FCM also promoted faster wound closure, with treated cultures approaching near-complete closure at later time points, while control cultures showed a more gradual decrease in wound area (Figure 2a, 2b). This migratory enhancement coincided with reduced proliferation, consistent with a “go-or-grow” phenotypic switch [13]. Immunofluorescence after 48 hours in FCM revealed a marked reduction in E-cadherin and upregulation of N-cadherin in both cell lines (Figure 2c). This cadherin switch is a canonical molecular feature of epithelial-to-mesenchymal transition [14], and these findings suggest that fibroblast-derived signals drive EMT-associated phenotypic plasticity in bladder cancer cells.

**Figure 2:**
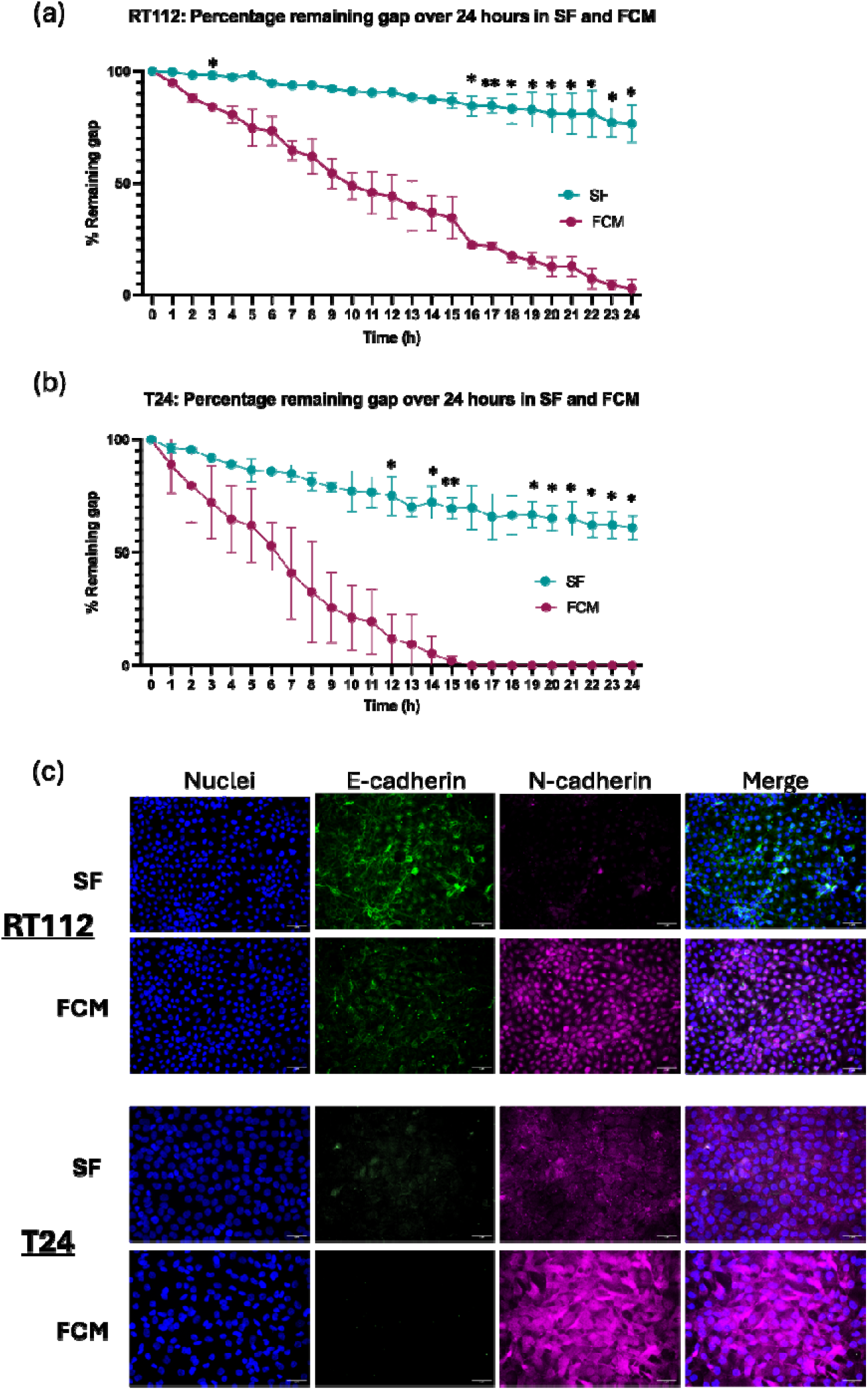
(a-b) Quantification of wound closure over time expressed as residual wound area (%) for RT112 (a) and T24 (b) via Leica MICA live imaging. FCM consistently increased the rate of wound closure compared with DMEM. Data are mean ± SD; n = 3; (c) Immunofluorescence for E-cadherin (green) and N-cadherin (magenta) with DAPI (blue) in RT112 and T24 after 48 h in DMEM or FCM. Immunofluorescence images taken at 40× magnification. Scale bars: 10µm.

### Bladder cancer cells induce fibroblast activation toward CAF-like states

To investigate whether cancer cells influence fibroblast activation, we first examined the effect of cancer cell-conditioned media on NFs. Specifically, we assessed the induction of CAF-like features by evaluating the expression of CAF markers, αSMA and FAP, after 48 hours of treatment. As shown in Figure 3a and 3b, exposure of NFs to RCM or TCM for 48 hours increased αSMA and FAP staining by immunofluorescence, particularly in TCM-treated cells.

**Figure 3:**
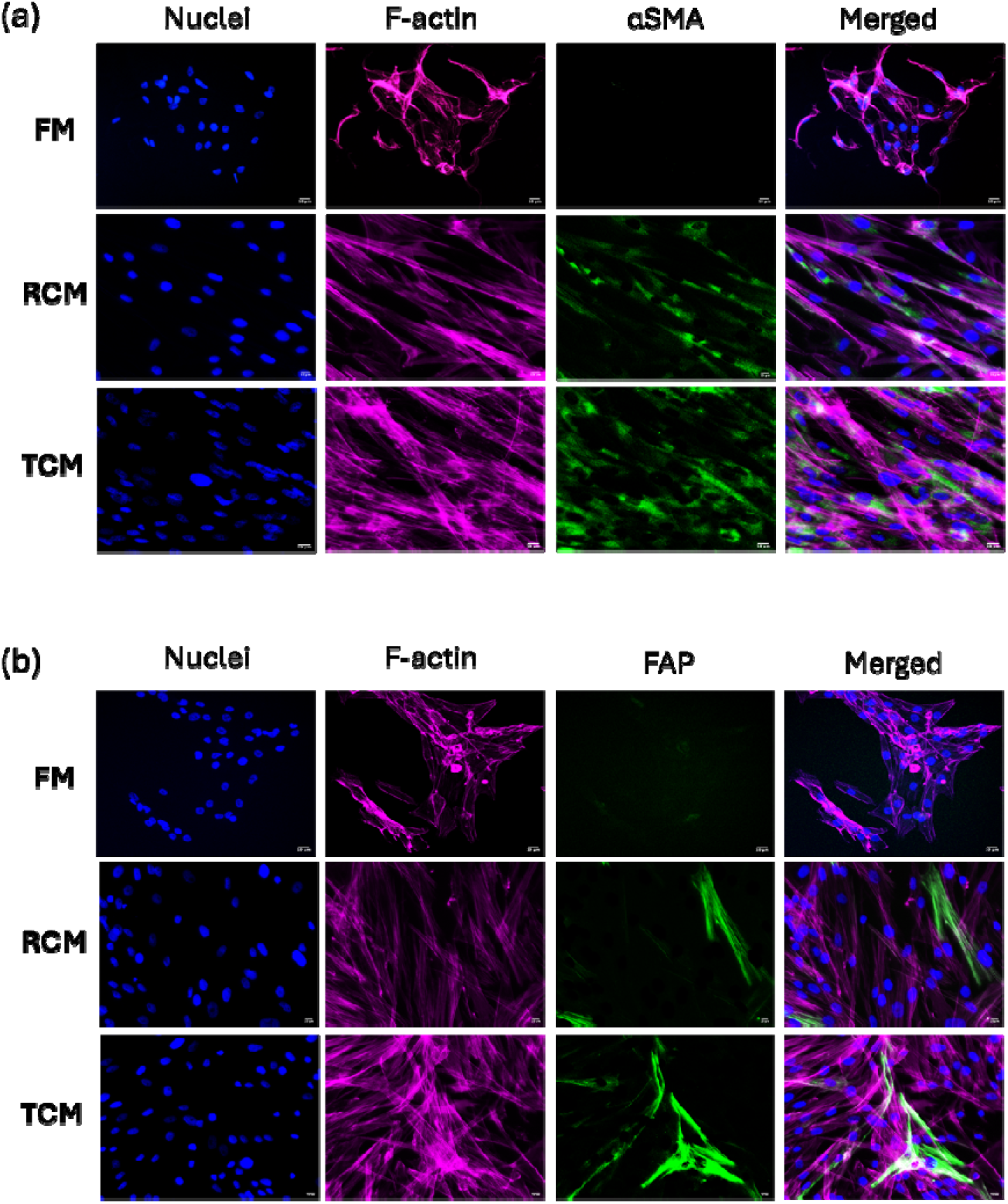
Detection of αSMA (a) and FAP (b) expression in NF after 48 hours of indirect co-culture with cancer-conditioned media. Immunofluorescence images taken at 20× magnification, showing positive green fluorescence indicating the presence of αSMA and FAP markers in NF. Scale bars: 10µm.

To quantify this in a direct contact model, NFs were co-cultured with RT112 or T24 and activation markers measured by flow cytometry, gating specifically on the NF population (Figure 4a, 4b, 4c). Basal αSMA and FAP levels were low in NF monoculture (1590.7 ± 210.4 and 3575.0 ± 552.2 MFI, respectively). Co-culture with RT112 drove αSMA to 2880.0 ± 522.7 MFI and FAP to 11,325.0 ± 823.0 MFI, a greater than three-fold induction for FAP, while T24 co-culture yielded comparable FAP induction (9823.3 ± 755.5 MFI) but more modest αSMA upregulation (2063.7 ± 257.6 MFI; Figure 4d, 4e). The divergence in αSMA induction between low-grade RT112 and high-grade T24 cells, despite comparable FAP responses, suggests grade-specific engagement of distinct CAF activation programmes.

**Figure 4:**
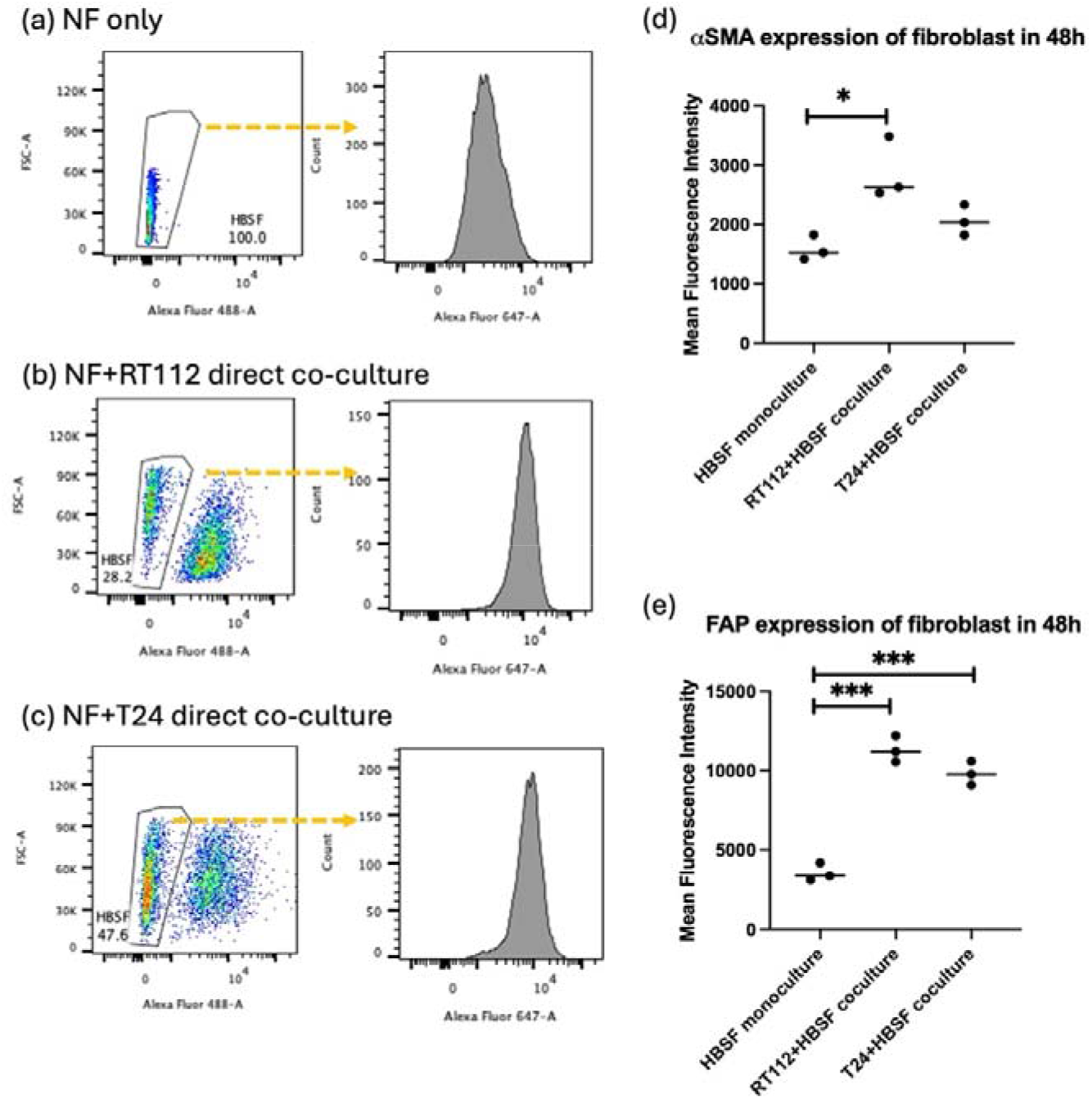
(a–c) Flow cytometry workflow and representative plots for (a) NF alone, (b) NF co-cultured with RT112, and (c) NF co-cultured with T24 for 48 h. Gates show total cells (FSC-A vs SSC-A), singlets (FSC-H vs FSC-A), and the NF population identified by Alexa Fluor 488 negative (pre-labelled cancer with CellTracker Green). The frequency of gated NF within each condition is indicated. Right-hand histograms display activation-marker staining in gated HBSF (αSMA/FAP; Alexa Fluor 647 channel). Quantification of mean fluorescence intensity (MFI) for (d) αSMA and (e) FAP in HBSF monoculture versus direct co-culture with RT112 or T24 (48 h). Both co-cultures significantly increased fibroblast activation marker expression, with the strongest induction observed in T24–HBSF. Bars show mean ± SD from n = 3.

Spatial organisation of tumour-stromal interactions, examined in 2D and 3D co-culture models, is shown in Supplementary Figure S1. In 2D co-culture, PanCK-positive tumour cells maintained discrete epithelial clusters while vimentin-positive fibroblasts adopted elongated morphologies and extended processes into surrounding tumour regions, forming a structured interface not observed in monoculture. T24 cells exhibited low-level vimentin expression consistent with their partial mesenchymal character. In 3D co-culture spheroids, fibroblasts were distributed throughout the spheroid mass rather than segregated peripherally, indicating close spatial integration between tumour and stromal compartments in both RT112 and T24 models. These spatial findings support the paracrine and contact-mediated crosstalk characterised in the preceding experiments.

### Cancer-fibroblast crosstalk attenuates MMC efficacy in a ratio-dependent manner

To determine whether stromal abundance modulates chemotherapy response, RT112 and T24 cells were treated with MMC (20 μM, 1 hour) in direct co-culture with NFs at increasing cancer-to-fibroblast ratios. Bulk viability was assessed by MTS assay, comparing observed co-culture survival against an expected value calculated from monoculture responses, assuming no interaction between populations. In RT112-NF co-culture, observed survival significantly exceeded the expected values across all fibroblast ratios (Figure 5a). In T24-NF co-culture, observed survival rates of 1:0.5 and 1:2 ratio co-culture were significantly higher than expected survival rates (Figure 5b). As bulk assays cannot distinguish whether this increase reflects true protection of cancer cells or simply the contribution of a less sensitive fibroblast population, we next performed flow cytometry to quantify cell death specifically within the cancer cell compartment. By separating CellTracker Red-labelled cancer cells from NFs and gating the dead cell population using the Zombie NIR viability kit, we found that the percentage of cell death induced by MMC in cancer cells was reduced in co-culture conditions compared with monoculture controls in both RT112 and T24 (Figure 5c and 5d). To further isolate the treatment-specific effect, we calculated ΔCell death% as the difference between treated and untreated within each ratio condition. This analysis showed that the net cytotoxic effect of MMC on cancer cells was progressively reduced in a presence of NFs (Figure 5e and 5f), confirming that fibroblast co-culture attenuates drug-induced killing rather than simply altering baseline viability. Together, these findings demonstrate that cancer-fibroblast interactions reduce MMC efficacy through active protection of tumour cells.

**Figure 5.**
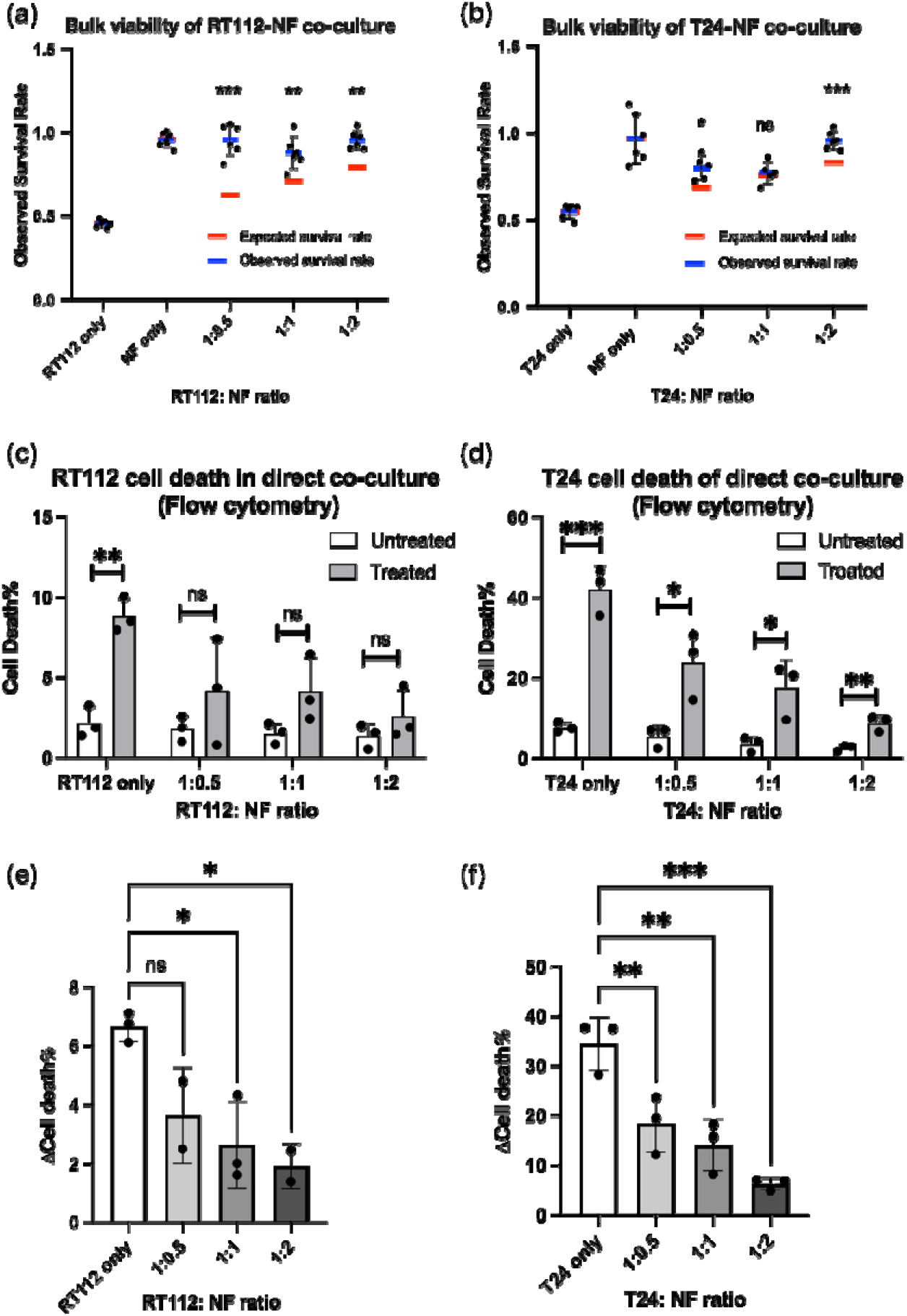
(a-b) MTS assay of RT112 (a) and T24 (b) cells treated with MMC in co-culture with NFs at increasing ratios. Observed survival in co-culture was compared with expected survival calculated from monoculture responses of cancer cells and fibroblasts, assuming no interaction between populations. (c-d) Flow cytometry-based quantification of cell death in cancer cell populations (RT112 (c) and T24 (d)) following direct co-culture with fibroblasts. Cancer cells were distinguished based on CellTracker Red labelling, enabling population-specific analysis. Co-culture significantly reduced MMC-induced cell death in cancer cells. (e–f) ΔCell death% in cancer cell populations, calculated as the difference cell death% between treated and untreated for RT112 (e) and T24 (f). Data are presented as mean ± SEM from 3 independent experiments.

### Fibroblast-rich tumours are associated with resistance signatures and poor clinical outcomes in bladder cancer

To determine whether the fibroblast-associated phenotypes observed in our experimental models were reflected in clinical data, we interrogated transcriptomic data from the US National Cancer Institute’s TCGA-BLCA database including samples from over 400 bladder cancer patients. A fibroblast-associated score was calculated for each tumour from a predefined fibroblast gene signature. The fibroblast-associated score exhibited a broad continuous distribution across the cohort (Supplementary Figure 2a), and tumours were stratified into fibroblast-low and fibroblast-high groups using the cohort median. To validate the biological relevance of this scoring approach, we examined its relationship with epithelial-mesenchymal transition (EMT) status. Fibroblast-associated scores showed a strong positive correlation with EMT scores across tumours (Spearman’s rho = 0.74, p < 0.0001; Supplementary Figure 2b), directly mirroring our in vitro finding that fibroblast-derived signals drive cadherin switching in cancer cells and confirming that the transcriptional fibroblast-enriched state maps onto a mesenchymal tumour phenotype in patients.

We then examined whether fibroblast-enriched tumours were associated with a transcriptional state consistent with therapy resistance. Fibroblast-high tumours showed increased expression of representative resistance-associated genes, including AXL, BCL2L1, MCL1, STAT3, CHEK1, and ERCC1 (Figure 6a), consistent with coordinated upregulation of adaptive survival signalling, anti-apoptotic pathways, and DNA damage response mechanisms. For example, AXL, a receptor tyrosine kinase linked to EMT and adaptive resistance, has also been reported to contribute to therapy resistance across multiple tumour types [15]. ERCC1 plays a central role in nucleotide excision repair and has been associated with resistance to DNA-damaging agents, including platinum-based therapies [16]. In addition, STAT3 is a key downstream effector of cytokine signalling, including CAF-derived IL-6, and has been implicated in promoting tumour cell survival and immune evasion in bladder cancer [17]. Consistent with these gene-level observations, fibroblast-high tumours displayed significantly elevated resistance-associated scores compared with fibroblast-low tumours (Figure 6b). Kaplan-Meier analysis demonstrated that patients with fibroblast-high tumours had significantly poorer overall survival than those with fibroblast-low tumours (Figure 6c). Together, these clinical cohort data mirror our in vitro findings that fibroblast-rich co-culture conditions attenuate therapy efficacy, and support a model in which tumour-fibroblast crosstalk promotes a transcriptionally defined, therapy-resistant state in bladder cancer.

**Figure 6.**
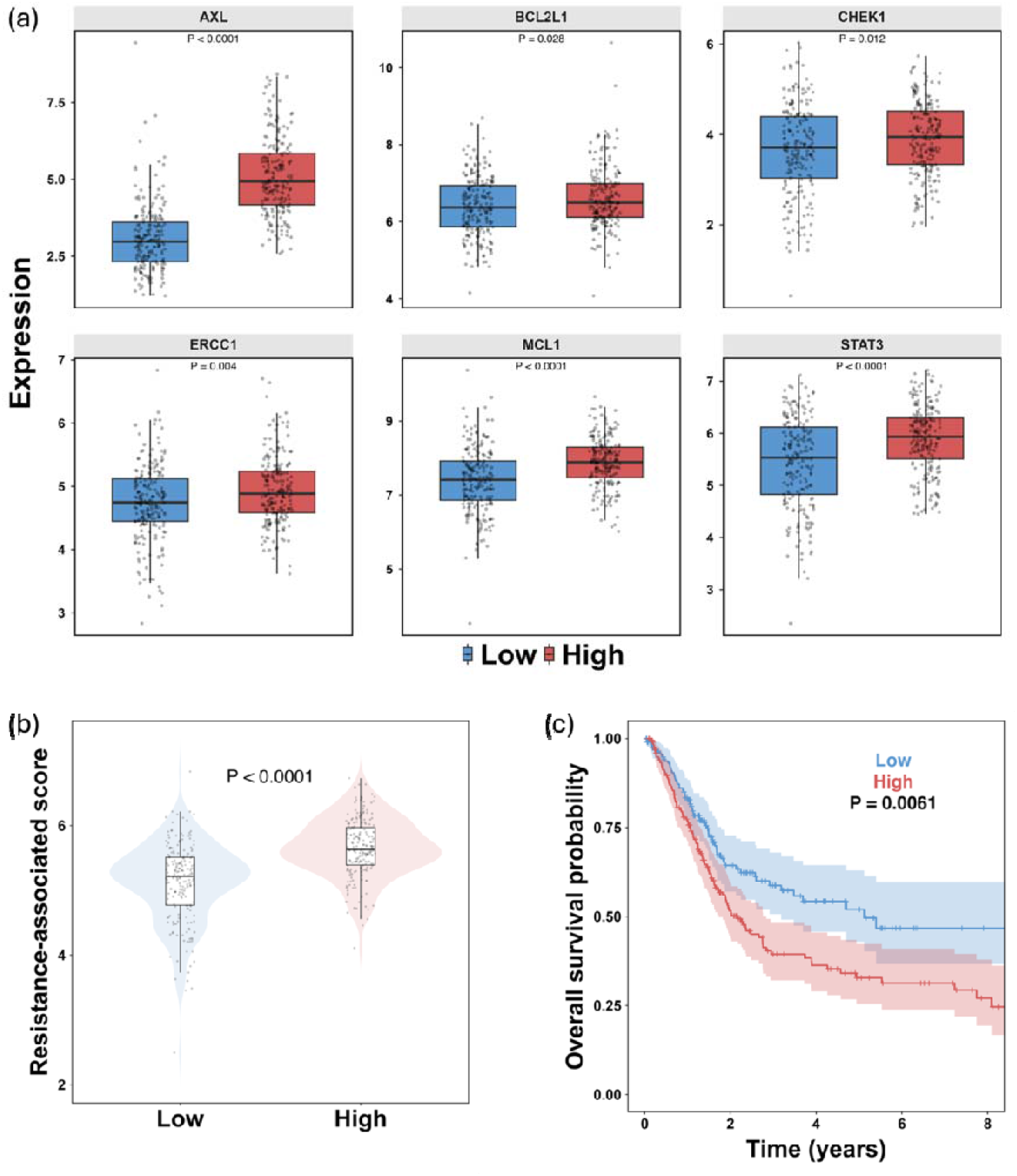
(a) Expression of representative resistance-associated genes (AXL, BCL2L1, CHEK1, ERCC1, MCL1 and STAT3) stratified by fibroblast score group in the TCGA-BLCA cohort. Fibroblast-high tumours showed increased expression of multiple resistance-related genes. (b) Comparison of resistance-associated signature scores between fibroblast-low and fibroblast-high tumours in the TCGA-BLCA cohort. Fibroblast-high tumours displayed significantly higher resistance scores than fibroblast-low tumours. (c) Kaplan Meier analysis of overall survival according to fibroblast score group in the TCGA-BLCA cohort. Patients with fibroblast-high tumours had significantly poorer overall survival than those with fibroblast-low tumours.

## Discussion

Tumour-stroma crosstalk is a recognised driver of cancer progression and therapy resistance, yet the specific role of normal fibroblasts, the important stromal population encountered by invading bladder cancer cells, has remained largely uncharacterised. This study demonstrates that this interaction is bidirectional, phenotypically consequential, and clinically reflected in patient transcriptomic data. Critically, prior work in bladder cancer has focused predominantly on already-activated CAFs [7]; this study, to the best of our knowledge, is the first to examine bidirectional tumour-NF crosstalk at the pre-activation stage, modelling the earliest microenvironmental encounter during lamina propria invasion.

FCM suppresses proliferation while enhancing migration, supporting a fibroblast-driven “go-or-grow” switch [13]. The responsible paracrine signals remain to be defined, but candidate mediators, including FGF, HGF, and CXCL12, engage RAS/MAPK and PI3K pathways that are known to promote EMT and invasive behaviour. The accompanying cadherin switch (E-to-N) provides molecular evidence that NF-derived signals promote a phenotypic state associated with invasive potential [14].

The differential αSMA and FAP induction between RT112 and T24 co-cultures is consistent with emerging evidence that bladder cancer CAFs comprise functionally distinct subpopulations, namely myofibroblastic CAFs (high αSMA) and inflammatory CAFs (high FAP/PDGFRα), which are differentially induced depending on tumour-derived signals [10]. In bladder cancer specifically, TGFβ is the primary driver of αSMA upregulation in stromal fibroblasts [18], suggesting that low-grade RT112 cells may preferentially engage TGFβ-mediated myofibroblastic activation. FAP expression in the stromal compartment carries independent prognostic significance in T1 NMIBC [19], underscoring the clinical relevance of NF-to-CAF conversion even at early invasion stages.

The fibroblast-mediated MMC resistance observed here likely reflects paracrine pro-survival signalling rather than physical drug barriers. CAF-secreted IL-6 and HGF activate STAT3 and AKT signalling to suppress apoptosis in tumour cells [7], a mechanism consistent with the elevated BCL2L1, MCL1, and STAT3 expression observed in fibroblast-high tumours from the TCGA. This transcriptional convergence between our in vitro MMC resistance data and patient gene expression profiles provides mutual validation across experimental and clinical dimensions.

Several limitations of the study warrant mention. We used a single commercially sourced NF line and simplified co-culture ratios that do not reflect patient stromal heterogeneity. The specific mediators of bidirectional crosstalk remain uncharacterised, and future work employing patient-derived organoid models will be required to establish causal mechanisms and validate fibroblast-associated signatures as predictive biomarkers of intravesical chemotherapy response.

In summary, our findings indicate that tumour-NF crosstalk in the bladder lamina propria drives EMT-like cancer cell reprogramming, fibroblast activation toward CAF-like states, and stromal protection against MMC. These findings are reflected in the transcriptional landscape from clinical tumour samples and the corresponding survival outcomes of patients. Targeting fibroblast-associated biomarkers in combination with intravesical MMC represents a potential strategy to overcome stromal-mediated resistance at the earliest stage of bladder cancer invasion.

## Materials and Methods

### Cell Culture

Bladder cancer cell lines RT112 (low-grade) and T24 (high-grade) and human bladder stromal fibroblasts (HBSF; referred to as NFs) were sourced from Dr Mark Linch’s Lab at the UCL Cancer Institute, ATCC, and ScienCell Research Laboratory, respectively. Cancer cells were cultured in DMEM (Gibco) with 10% FBS (Fisher Scientific) and 1% Penicillin/Streptomycin (Fisher Scientific). NFs were maintained in Fibroblast Medium (FM; ScienCell #2301). All cells were maintained at 37°C, 5% CO_2_ and passaged at 80% to 90% confluence using Accutase (Merck).

### Conditioned Medium Preparation

Serum-free DMEM was used to eliminate FBS-associated variability. RT112, T24, and NFs were seeded in T150 flasks respectively. After 24 hours, media were replaced with serum-free DMEM containing 1% Penicillin/Streptomycin. After 48 hours of incubation, conditioned media RCM, TCM, and FCM were harvested, centrifuged to remove debris, filtered through 0.2 µm filters (Fisher Scientific), and stored at 4°C for subsequent experiments.

### MTS Cell Proliferation Assay

For conditioned media experiments, 3,000 cancer cells or NFs per well were seeded in 96-well plates, treated with conditioned or serum-free media after 24 hours, and assessed by MTS assay (Promega) at 48 and 96 hours. For MMC experiments, RT112 or T24 cells were co-cultured with NFs at 1:0, 1:1, and 1:2 ratios (6,000 total cells/well). After 24 hours, cells were treated with MMC (Selleck Chemicals, 20 μM, 1 hour), washed with PBS (Fisher Scientific), and incubated in fresh medium for 72 hours before the MTS assay. To determine whether co-culture survival exceeded what would be expected from each cell population independently, an expected survival value was calculated for each ratio as a weighted average of monoculture MMC responses: Expected survival (%) = [f_cancer × Survival_cancer] + [f_NF × Survival_NF], where f_cancer and f_NF are the fractional contributions of each population at the given seeding ratio, and Survival_cancer and Survival_NF are the MTS-measured survival percentages of cancer cells and NFs in MMC-treated monoculture, respectively. Observed survival significantly exceeding expected survival indicates active stromal protection beyond the additive contributions of each population.

### Wound Healing Assay - Cell Migration

RT112 (2.5 × 10□) and T24 (2.0 × 10□) cells were seeded to confluence in 24-well plates. A wound gap was created using Ibidi Culture-Insert 2-Well inserts, which were removed after 24 hours. Medium was replaced with either serum-free DMEM or FCM at the time of insert removal (t = 0). Images were captured hourly for 24 hours using a Leica MICA microscope and residual wound gap (%) was measured using LAS X software, normalised to the t = 0.

### Immunofluorescence

For indirect co-culture, NFs or cancer cells (3 × 10□/well) were seeded on Ibidi µ-slides, treated with conditioned media for 48 hours, then fixed in 4% formaldehyde, permeabilised with 0.2% Triton X-100 (Fisher Scientific), and blocked with 5% normal goat serum/1% BSA. NFs were stained with anti-FAP (rabbit monoclonal, Cell Signalling Technology, CST, #66562) and anti-αSMA (rabbit monoclonal, CST, #19245) at 1:100, followed by anti-rabbit Alexa Fluor 488 and Alexa Fluor 555 Phalloidin (Invitrogen). Cancer cells were stained with anti-E-cadherin (mouse monoclonal, Invitrogen, #33-4000; 10 μg/ml) and anti-N-cadherin (rabbit polyclonal, Invitrogen, #PA5-19486; 1 μg/ml), followed by appropriate fluorescently labelled secondary antibodies. Nuclei were counterstained with DAPI (Fisher Scientific). For 2D direct co-culture and 3D spheroid immunofluorescence, RT112 or T24 cells and NFs were co-seeded at a 1:1 ratio on µ-slide 8-well chamber slides (Ibidi) and cultured for 48 hours. Cells were fixed via 4% formaldehyde (FA) (Fisher Scientific) and stained for pan-cytokeratin (mouse monoclonal, CST, #67306) and vimentin (rabbit monoclonal, CST, #5741) at 1:100, with DAPI counterstain. For 3D spheroid generation, RT112 or T24 cells and NFs were mixed at a 1:1 ratio and seeded in ultra-low attachment 96-well round-bottom plates (Fisher Scientific) at a total density of 3,000 cells per well. Spheroids were allowed to form over 72 hours and imaged by Leica SP8 confocal microscopy following fixation and immunofluorescence staining for PanCK and vimentin at 1 in 100 dilution as well.

### Flow Cytometry

Prior to co-culture, RT112 or T24 cancer cells were labelled with CellTracker Green CMFDA (Thermo Fisher Scientific, C7025; 10 μM in serum-free DMEM for 30 minutes at 37°C) to enable downstream discrimination from NFs by flow cytometry. Labelled cancer cells were then co-seeded with NFs in T75 flasks at a 1:1 ratio for 48 hours. Cells were detached with Accutase and fixed with 4% FA. After permeabilisation with 0.2% Triton X-100 and blocking with 5% normal goat serum, cells were stained with αSMA or FAP primary antibodies (1:100) for 45 minutes. Secondary antibody (anti-rabbit Alexa Fluor 488) was applied for 30 minutes at 1:500. After washing, cells were resuspended in 1% BSA and analysed using a BD LSR Fortessa flow cytometer. Data were processed using FlowJo software.

To enable population-specific cell death analysis, cancer cells (RT112 or T24) were labelled with CellTracker Red CMTPX (Thermo Fisher Scientific, C34552; 5 μM in serum-free DMEM for 30 minutes at 37°C) and NFs were labelled with CellTracker Green CMFDA (Thermo Fisher Scientific, C7025; 10 μM in serum-free DMEM for 30 minutes at 37°C) prior to co-culture. Red labelled cancer cells were seeded with green labelled NFs at 1:0, 1:1, and 1:2 ratios in 6 well plates (total 300,000 cells/well). After 24 hours, cells were treated with MMC (20 μM, 1 hour), washed with PBS, and incubated in fresh medium for 72 hours. Cells were then detached with Accutase, pelleted, and resuspended in PBS. Cell viability was assessed using Zombie NIR Fixable Viability Kit (BioLegend, #423105; 1:1000 dilution in PBS for 15 minutes at room temperature, protected from light). After washing, cells were resuspended in 1% BSA/PBS and acquired on a BD LSRFortessa. Cancer cells were gated as CellTracker Red-positive; Zombie NIR-positive cells within this gate were scored as dead. Cell death% was defined as the percentage of Zombie NIR-positive cells within the CellTracker Red-positive population. ΔCell death% was calculated as: (cell death% in MMC-treated condition) − (cell death% in matched untreated condition) for each co-culture ratio, isolating the treatment-specific cytotoxic effect from baseline viability differences. Data were processed in FlowJo.

### Public transcriptomic cohort analysis

TCGA-BLCA gene expression (log2-TPM) and clinical data were obtained from the GDC data portal. A fibroblast-associated score (mean expression of FAP, ACTA2, TAGLN, PDGFRB, COL1A1, COL1A2, FN1, POSTN, THY1, CXCL12, TGFB1) and a resistance-associated score (mean expression of AXL, BCL2L1, MCL1, XIAP, BIRC5, STAT3, YAP1, ITGB1, CHEK1, ERCC1) were computed per sample. Tumours were stratified into fibroblast-high/low groups at the cohort median. An EMT score was calculated as the mean expression of a predefined mesenchymal gene signature (VIM, FN1, CDH2, TWIST1, SNAI1, SNAI2, ZEB1, ZEB2) minus the mean expression of an epithelial gene signature (CDH1, EPCAM, KRT8, KRT18, KRT19, MUC1), consistent with established approaches for transcriptomic EMT scoring in bladder cancer. Statistical analyses used Wilcoxon rank-sum tests, Spearman correlation, and Kaplan-Meier log-rank tests in R (v4.5.2).

### Statistical Analysis

In vitro data were analysed by unpaired two-tailed t-test. Data are presented as mean ± SEM. p ≤ 0.05 was considered significant. GraphPad Prism v10.3.0 was used for graphing. *p ≤ 0.05; **p ≤ 0.01; ***p ≤ 0.001; ****p ≤ 0.0001.

## Declaration

Declarations of interest: JLR and ES have share options in AtoCap Ltd, who focus is not related to the present manuscript.

## CRediT authorship contribution statement

JG: Conceptualization; Methodology; Investigation; Funding acquisition; Supervision; Writing original draft; Review & editing; CXJ: Investigation; Methodology; CR: Investigation; Methodology; JH: Review & editing; ES: Funding acquisition; Supervision; Review & editing. RTB: Funding acquisition; Supervision; Review & editing. JLR: Funding acquisition; Supervision; Review & editing.

## References

[1] Richters A, Aben KKH, Kiemeney LALM. The global burden of urinary bladder cancer: an update. World J Urol 2020;38:1895–904. 10.1007/s00345-019-02984-4.

[2] Cox E, Saramago P, Kelly J, Porta N, Hall E, Tan WS, et al. Effects of Bladder Cancer on UK Healthcare Costs and Patient Health-Related Quality of Life: Evidence From the BOXIT Trial. Clin Genitourin Cancer 2020;18:e418–42. 10.1016/j.clgc.2019.12.004.

[3] Grabe-Heyne K, Henne C, Mariappan P, Geiges G, Pöhlmann J, Pollock RF. Intermediate and high-risk non-muscle-invasive bladder cancer: an overview of epidemiology, burden, and unmet needs. Front Oncol 2023;13:1170124. 10.3389/fonc.2023.1170124.

[4] Scilipoti P, Ślusarczyk A, De Angelis M, Soria F, Pradere B, Krajewski W, et al. The Role of Mitomycin C in Intermediate-risk Non–muscle-invasive Bladder Cancer: A Systematic Review and Meta-analysis. Eur Urol Oncol 2024;7:1293–302. 10.1016/j.euo.2024.06.005.

[5] Claps F, Pavan N, Ongaro L, Tierno D, Grassi G, Trombetta C, et al. BCG-Unresponsive Non-Muscle-Invasive Bladder Cancer: Current Treatment Landscape and Novel Emerging Molecular Targets. Int J Mol Sci 2023;24:12596. 10.3390/ijms241612596.

[6] Lee Y-C, Lam H-M, Rosser C, Theodorescu D, Parks WC, Chan KS. The dynamic roles of the bladder tumour microenvironment. Nat Rev Urol 2022;19:515–33. 10.1038/s41585-022-00608-y.

[7] Huang L, Xie Q, Deng J, Wei W-F. The role of cancer-associated fibroblasts in bladder cancer progression. Heliyon 2023;9:e19802. 10.1016/j.heliyon.2023.e19802.

[8] Barbazan J, Pérez-González C, Gómez-González M, Dedenon M, Richon S, Latorre E, et al. Cancer-associated fibroblasts actively compress cancer cells and modulate mechanotransduction. Nat Commun 2023;14:6966. 10.1038/s41467-023-42382-4.

[9] Mao X, Xu J, Wang W, Liang C, Hua J, Liu J, et al. Crosstalk between cancer-associated fibroblasts and immune cells in the tumor microenvironment: new findings and future perspectives. Mol Cancer 2021;20:131. 10.1186/s12943-021-01428-1.

[10] Burley A, Rullan A, Wilkins A. A review of the biology and therapeutic implications of cancer-associated fibroblasts (CAFs) in muscle-invasive bladder cancer. Front Oncol 2022;12:1000888. 10.3389/fonc.2022.1000888.

[11] Greenwell JC, Torres-Gonzalez E, Ritzenthaler JD, Roman J. Fibroblast-derived conditioned media promotes lung cancer progression. Am J Med Sci 2023;365:189–97. 10.1016/j.amjms.2022.08.019.

[12] Koh B, Jeon H, Kim D, Kang D, Kim K. Effect of fibroblast co⍰culture on the proliferation, viability and drug response of colon cancer cells. Oncol Lett 2018. 10.3892/ol.2018.9836.

[13] Crossley RM, Painter KJ, Lorenzi T, Maini PK, Baker RE. Phenotypic switching mechanisms determine the structure of cell migration into extracellular matrix under the ‘go-or-grow’ hypothesis. Math Biosci 2024;374:109240. 10.1016/j.mbs.2024.109240.

[14] Loh C-Y, Chai J, Tang T, Wong W, Sethi G, Shanmugam M, et al. The E-Cadherin and N-Cadherin Switch in Epithelial-to-Mesenchymal Transition: Signaling, Therapeutic Implications, and Challenges. Cells 2019;8:1118. 10.3390/cells8101118.

[15] Wu F, Li J, Jang C, Wang J, Xiong J. The role of Axl in drug resistance and epithelial-to-mesenchymal transition of non-small cell lung carcinoma. Int J Clin Exp Pathol 2014.

[16] Urun Y, Leow JJ, Fay AP, Albiges L, Choueiri TK, Bellmunt J. ERCC1 as a prognostic factor for survival in patients with advanced urothelial cancer treated with platinum based chemotherapy: A systematic review and meta-analysis. Crit Rev Oncol Hematol 2017;120:120–6. 10.1016/j.critrevonc.2017.10.012.

[17] Mirzaei S, Gholami MH, Mahabady MK, Nabavi N, Zabolian A, Banihashemi SM, et al. Pre-clinical investigation of STAT3 pathway in bladder cancer: Paving the way for clinical translation. Biomed Pharmacother 2021;133:111077. 10.1016/j.biopha.2020.111077.

[18] Xu Y, Li W, Lin S, Liu B, Wu P, Li L. Fibroblast diversity and plasticity in the tumor microenvironment: roles in immunity and relevant therapies. Cell Commun Signal 2023;21:234. 10.1186/s12964-023-01204-2.

[19] Muilwijk T, Baekelandt L, Akand M, Daelemans S, Marien K, Waumans Y, et al. Fibroblast Activation Protein-α and the Immune Landscape: Unraveling T1 Non–muscle-invasive Bladder Cancer Progression. Eur Urol Open Sci 2024;66:67–74. 10.1016/j.euros.2024.06.011.

