## Supplemental data for "Bidirectional Crosstalk Between Bladder Cancer Cells and Normal Fibroblasts Drives Phenotypic Reprogramming and Mitomycin C Resistance"

### Supplementary Results

Direct 2D and 3D co-culture models reveal distinct organisation of tumour-fibroblast interactions

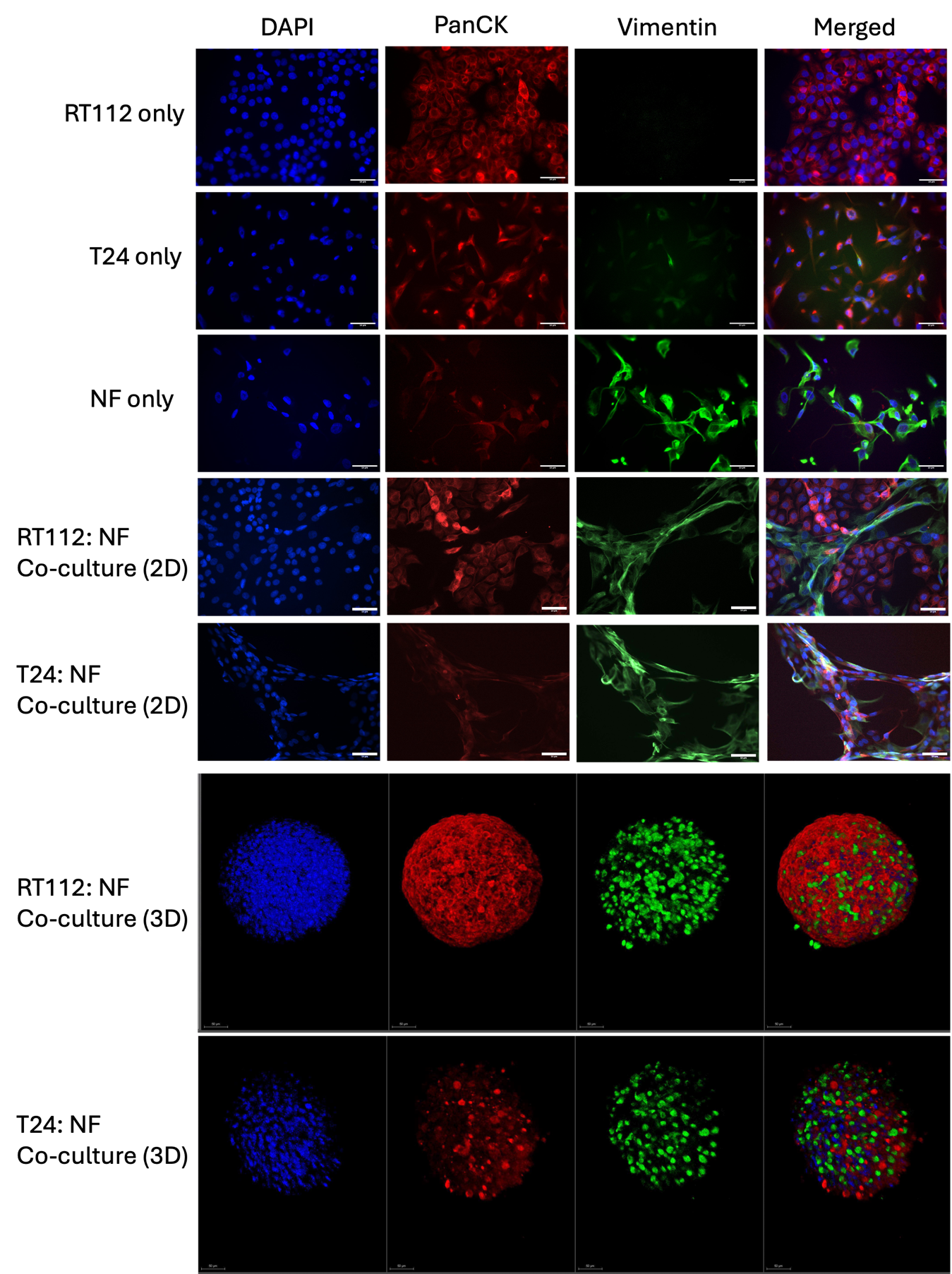

Sup-Figure 1: Immunofluorescence staining of bladder cancer cells (RT112, T24) and NF. PanCK (red) marks epithelial cells, vimentin (green) marks stromal fibroblasts, and nuclei are stained with DAPI (blue). Scale bars: 50 µm

Tumour-fibroblast spatial organisation was examined using direct co-culture of bladder cancer cells with NF, followed by immunofluorescence staining for epithelial (PanCK) and mesenchymal (vimentin) markers. In direct 2D co-culture, tumour and stromal cells formed a structured interface. PanCK-positive tumour cells maintained epithelial clusters, whereas vimentin-positive fibroblasts adopted elongated morphologies and extended processes surrounding tumour cell regions, creating an organised tumour-stromal architecture not observed in monoculture conditions. Additionally, T24 cells show low-level vimentin staining consistent with partial mesenchymal features of high-grade bladder cancer.

To further investigate tumour-stromal organisation in a three-dimensional context, tumour-fibroblast spheroids (co-culture at 1 in 1 ratio) were generated and analysed by Leica SP8 confocal microscopy. In these spheroids, PanCK-positive tumour cells formed the main spheroid structure, while vimentin-positive fibroblasts were distributed throughout the spheroid mass, indicating close spatial association between tumour and stromal components. Similar organisational patterns were observed in both RT112 and T24 co-culture spheroids.

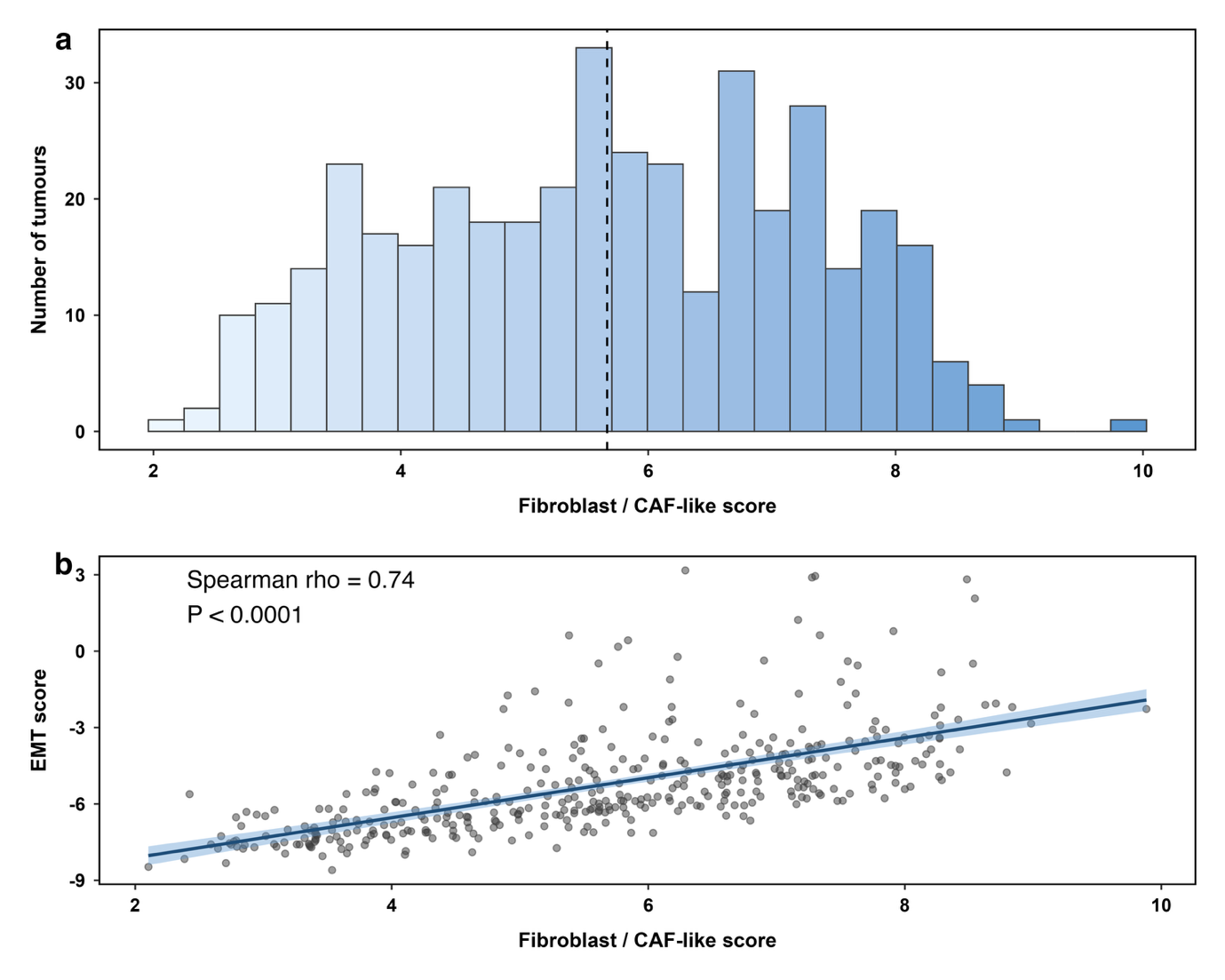

Sup-Figure 2. (a) Distribution of fibroblast-associated scores across TCGA-BLCA tumours. The dashed line indicates the cohort median used to stratify tumours into fibroblast-low and fibroblast-high groups. (b) Correlation between fibroblast-associated score and EMT score across tumours. EMT score was calculated as the difference between mesenchymal and epithelial gene expression signatures. Each point represents an individual tumour sample. A strong positive correlation was observed (Spearman’s rho = 0.74, P < 0.0001), indicating that fibroblast-enriched tumours are associated with an EMT-like transcriptional state.

To further characterise the biological features associated with fibroblast enrichment, we examined the relationship between fibroblast-associated scores and epithelial-mesenchymal transition (EMT) programmes in the TCGA-BLCA cohort. Fibroblast-associated scores displayed a broad distribution across tumours, with a median value used for downstream stratification (Sup-Figure 2a). Notably, fibroblast-associated scores showed a strong positive correlation with EMT scores (Spearman’s rho = 0.74, P < 0.0001; Sup-Figure 2b), indicating that fibroblast-enriched tumours are associated with a mesenchymal-like transcriptional state.

These findings are consistent with our in vitro observations that fibroblast-derived signals promote an EMT-like phenotype in bladder cancer cells, and suggest that tumour-fibroblast interactions contribute to the emergence of migratory and potentially therapy-resistant tumour states in patients.
